# Deletion of Autophagy gene *ATG1* and Glyoxylate cycle Regulator encoding gene *UCC1* together leads to Synthetic Growth Defects and Sensitivity to Genotoxic agents in *Saccharomyces cerevisiae*

**DOI:** 10.1101/2020.03.03.974568

**Authors:** Monika Pandita, Vishali Sharma, Meenu Sharma, Heena Shoket, Sadia Parvez, Prabhat Kumar, Narendra K Bairwa

**Author notes:** contributed equally (Monika Pandita, Vishali Sharma, Meenu Sharma). To whom correspondence may be addressed, /285634.

## Abstract

Atg1 of *S. cerevisiae* is a key component of autophagy encoded by *ATG1* gene, involved in the process of degradation of cytosolic components through autophagy. *UCC1*, an F-box encoding gene is involved in the negative regulation of glyoxylate pathway via degradation of Cit2 enzyme by ubiquitin proteasome system. We investigated the genetic interaction between *ATG1* and *UCC1* using the gene deletion approach. The *atg1Δucc1Δ* cells showed the synthetic growth defects with abnormal budding and sensitivity to genotoxic and oxidative stress agents. Based on the observations, we report that *ATG1* and *UCC1* interact genetically to regulate the cell growth fitness and function in parallel pathway in cellular response to the genotoxic stress agents. The present investigation also revealed the cross talks among autophagy, ubiquitin proteasome system, and glyoxylate pathways.

## Introduction

Autophagy is a conserved degradative process for the degradation of cytoplasmic organelles by the hydrolytic enzymes in the vacuole when exposed to starvation conditions (Iwama and Ohsumi 2019). The process includes the formation of double-membrane structure, called auto-phagosome, followed by fusion with vacuole lumen and disintegration of the autophagic body by hydrolytic enzymes for release of simple molecules in the cytoplasm, for reuse in the various cellular processes (Thumm 2000). Autophagy plays critical role in various physiological processes and is a preferred target for precision therapeutics and drug development. There are 32 different autophagy related genes (*ATG* genes) that have been identified by genetic screening in yeast and many of these genes are conserved across insects, plants, annelids, and mammals. The *ATG1* or *YGL180W* gene in *S. cerevisiae* is a serine-threonine kinase molecule that is evolutionarily conserved. The Atg1 forms the complex with other initiator units of autophagy including Atg13, Atg17, Atg29, and Atg31. This molecular complex is involved in the elimination of damaged organelles or misfit proteins with altered structural folding to preserve homeostasis (Davies *et al.* 2015). Atg1 has been reported in the cell cycle progression from G2/M phase to G1 during nitrogen deprived conditions (Matsui *et al.* 2013). The mutation in *ATG1* leads to negative influence on the autophagy cycle (Papinski and Kraft 2016).

*UCC1* or *YLR224W* gene of *S. cerevisiae* encodes for F-box motif containing protein, Ucc1, which forms the complex with the SCF E3 ligase of ubiquitin proteasomal system (UPS) to recruit the Cit2, citrate synthase of the glyoxylate cycle that aids in the utilization of alternately available, non-fermentable carbon sources (Nakatsukasa *et al.* 2015) for ubiquitination and degradation by the 26S proteasome. Ucc1 over expression leads to induced n-butanol tolerance whereas deletion mutant showed butanol sensitivity, implicating the involvement of *UCC1* in C3 and C4 alcohol utilization (GonzÁlez-Ramos *et al.* 2013). In this study, we investigated the genetic interaction between *ATG1* and *UCC1* and their role in the growth fitness and response to genotoxic (MMS, HU) and oxidative stress (H_2_O_2_). Here we report that the absence of both the gene *ATG1* and *UCC1* together leads to synthetic growth defects and sensitivity to genotoxic and oxidative stress.

## Methods

### Strains, Plasmids and Growth media

*Saccharomyces cerevisiae* laboratory strain BY4741 (*MATa his3Δ1 leu2Δ0 met15Δ0 ura3Δ0*) was used for the construction of deletion mutants of *ATG1* and *UCC1*. Plasmids, pFA6A-*KanMX6*, and pFA6A-His3MX6 were used for generation of PCR product containing the selectable marker cassette with 40 bp homology region to the flanking region of the target ORF sequence. For transformation of the PCR product into the yeast cell and selection, method mentioned in the (Gietz *et al.* 1995; Longtine *et al.* 1998) was followed. The 200 μg/ml of G418 was used for *KanMX6* marker cassette selection. The list of primers used for the ORF deletion and yeast strains are mentioned in the **Table 1, Table 2**.

**Table 1:**
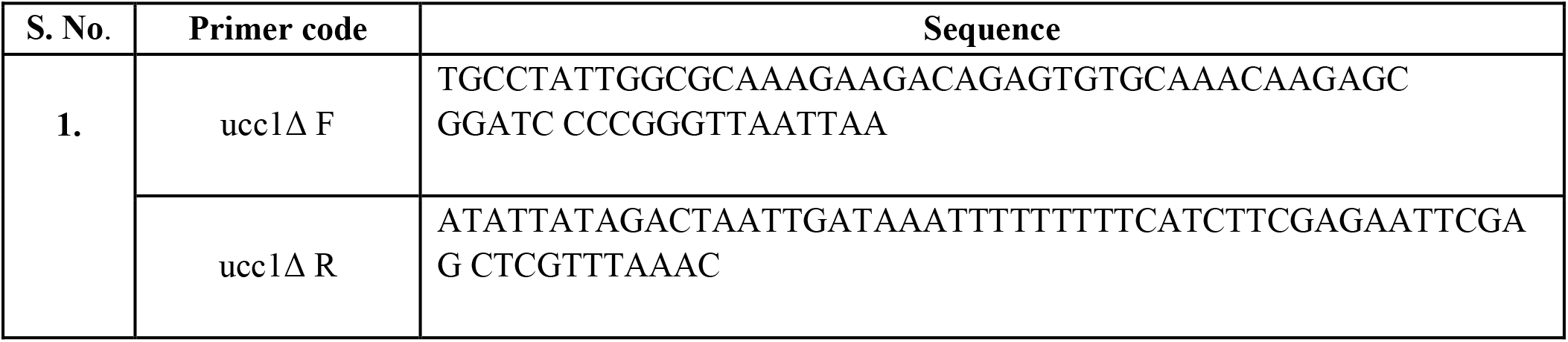
List of primers used in the study for construction of deletion strains.

**Table 2:** Yeast strains used in the study and their genotypes.

| S. No. | Strain | Genotype | Source |
| --- | --- | --- | --- |
| 1. | BY4741 | *MATa his3Δ1 leu2Δ0 met15Δ0*<br>*ura3Δ0* | Dr. Deepak Sharma,<br>IMTECH |
| 2. | MP1 | *ucc1Δ::KanMX* | This study |
| 3. | MNK1 | *atg1Δ::His3MX* | Dr. Ravi Manjithya,<br>JNCASR, India |
| 4. | MNK2 | *ucc1Δ::KanMX atg1Δ::His3MX* | This study |

### Growth assay

For comparative growth analysis of the strains on solid media (YPD+agar), single colony of each strain was streaked and incubated at the 30°C for 36-48 hrs and growth was assessed and photographed. To compare the growth kinetics, single colony of each strain was inoculated into 25 ml of YPD broth and incubated overnight at 30°C. Next day each culture was transferred to autoclaved 50 ml YPD media in 1:20 dilution ratio and an optical density (OD) at 600nM was taken to adjust the each culture by OD. The initial OD was taken as starting time point and culture was allowed to grow at 30°C. After every 2 hrs OD was taken and a growth curve was plotted as time versus OD to compare the growth between strains.

### Microscopy

For morphological assessment each strain was grown till log phase and imaged using Leica DM3000 microscope with 100X magnification using phase filters. For analysis of the chitin distribution of in the cell wall, each strain was grown to log phase and stained with the **c**alcofluor white fluorescent dye as mentioned in (Blanco *et al.* 2012; Preechasuth *et al.* 2015). Briefly, cells were grown overnight at 30⁰ C. After incubation for 12-14 hours, cells were transferred in fresh YPD in a 1:10 ratio. Cells were grown till log phase following which 1 ml of cells was collected by centrifugation. Cells were washed with 1X PBS twice and stained with Calcofluor white fluorescent dye (50μg/ml) and observed under Leica DM3000 fluorescence microscope at 100X magnification. To analyse the nucleus status in the strains, yeast strains were grown at 30⁰C till log phase in YPD medium Cells were collected by centrifugation at 2000 rpm for 3 minutes and washed with distilled water, then suspended in 1X PBS. Cells were fixed with 70% ethanol, washed in 1 X PBS, and centrifuged for 1 minute at 2500 rpm. Cell were stained by adding 2.5μg/ml of final concentration (1mg/ml DAPI stock) DAPI and incubated at room temperature for 3-5 minutes. Cells were analysed using Leica DM3000 fluorescence microscope under UV light at 100X magnification.

### Spot assay for cellular growth response to genotoxic and oxidative stress

For analysis of the cellular growth response to genotoxic (0.035% MMS, 200mM hydroxyurea) and oxidative stress (4mM Hydrogen peroxide), cells were grown up to log phase in YPD at 30⁰C, harvested, washed and tenfold serially diluted. From each dilution, 3 μl of sample was spotted on YPD agar plates and plates containing stress agents. Plates were incubated for 24-36 hrs at 30⁰C and photographed.

## Results

### Loss of *ATG1* and *UCC1* together leads to synthetic growth defects

*ATG1* regulates the process of autophagy in *S. cerevisiae* to help cells survive stress conditions, particularly nutrient non-availability or exposure to infectious agents and thus by disintegration of cellular components into recyclable simple molecules, the process becomes “pro-survival”(Gump and Thorburn 2011). The *atg1* null mutant showed a decrease in viability and a decline to chemical based resistance (Tsukada and Ohsumi 1993) whereas absence of *ucc1* doesn’t hold a major influence on the cell viability (Nakatsukasa *et al.* 2015). We investigated the genetic interaction between *ATG1* and *UCC1* and impact on the growth fitness. We observed that double mutant *atg1Δucc1Δ* showed slow growth phenotype when compared with the WT, *atg1Δ, and ucc1Δ* cells (**Fig 1a, b).** Morphological assessment of the WT, *atg1Δ, and ucc1Δ* and *atg1Δucc1Δ* cells by microscopy indicated abnormal bud formation in *atg1Δucc1Δ* with clumps formation (**Fig.1c)**. The data indicates that *ATG1* and *UCC1* together are essential for cell growth fitness.

**Figure 1:**
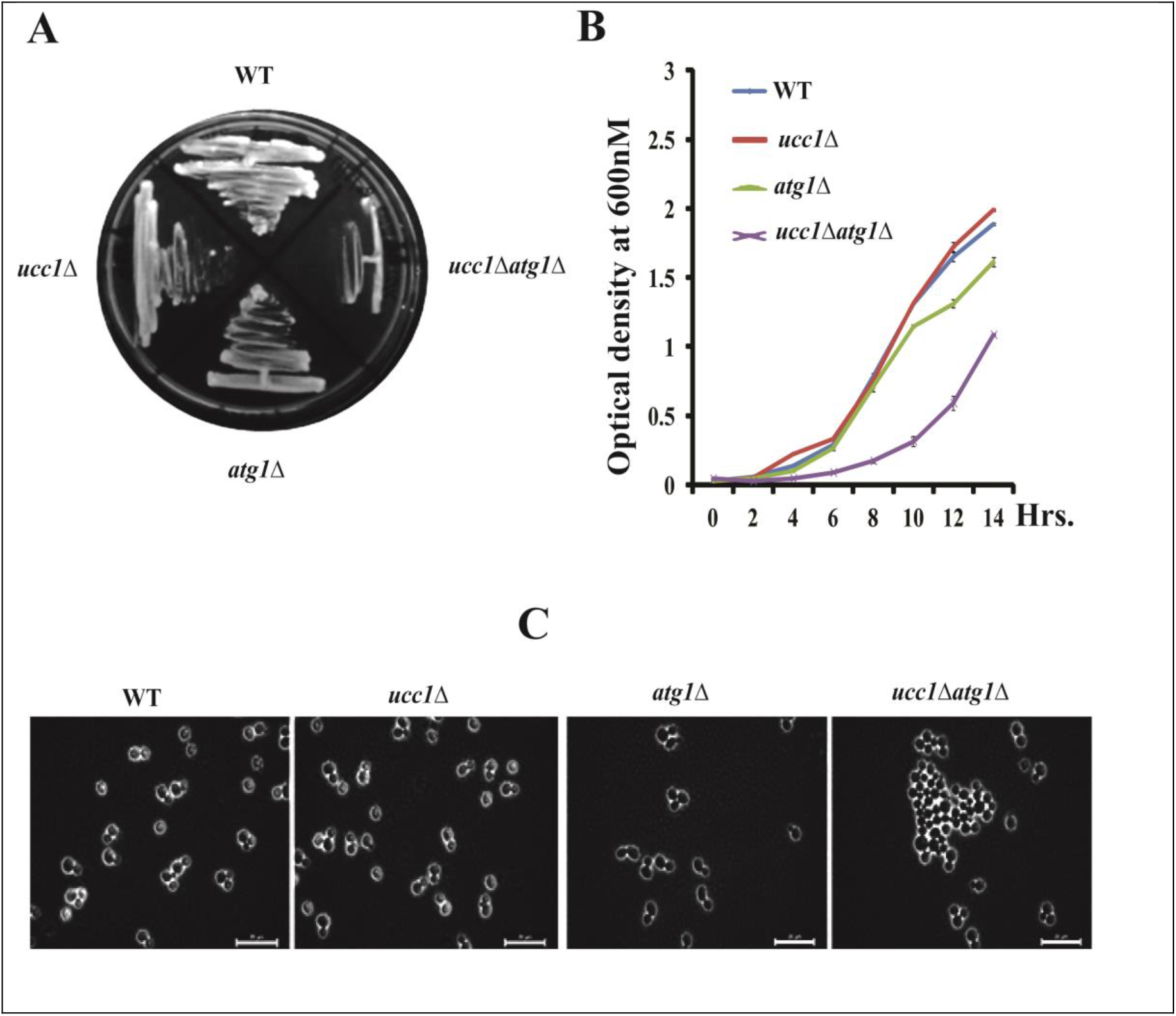
Comparative assessment of growth and morphological characteristics of WT, *ucc1* Δ,*atg1Δ, ucc1* Δ *atg1* Δ strains. **A** Comparative growth on solid medium after streaking single colony and incubation for 24-36 hrs at 30°C **B.** Growth kinetics of WT, *ucc1*Δ, *atg1*Δ and *ucc1*Δ*atg1*Δ in YPD liquid medium. Cells were collected every 2 hours interval and cellular growth was analysed by measurement of optical density (OD) at 600nm using TOSHVIN UV-1800 SHIMADZU. The graph plotted represents the average of three independent replicates. The error bars represent standard deviation (SD) for each data set. **C.** Phase contrast imaging of log phase grown cells at 100X magnification using Leica DM3000 microscope.

### Loss of *ATG1* and *UCC1* together leads to altered chitin level

Cell wall is composed of linear chains of chitin molecules that border the peripheral region of cell. The analysis of the chitin status in the WT, *atg1Δ*, *ucc1Δ* and *atg1Δucc1Δ* by staining with Calcofluor white fluorescent stain indicated no major change in the chitin distribution. The WT cells showed the staining only in the bud region, whereas the *atg1Δ, and ucc1Δ* and *atg1Δucc1Δ* mutants exhibited high concentration of chitin at bud adjacent region (**Figure 2)**.

**Figure 2:**
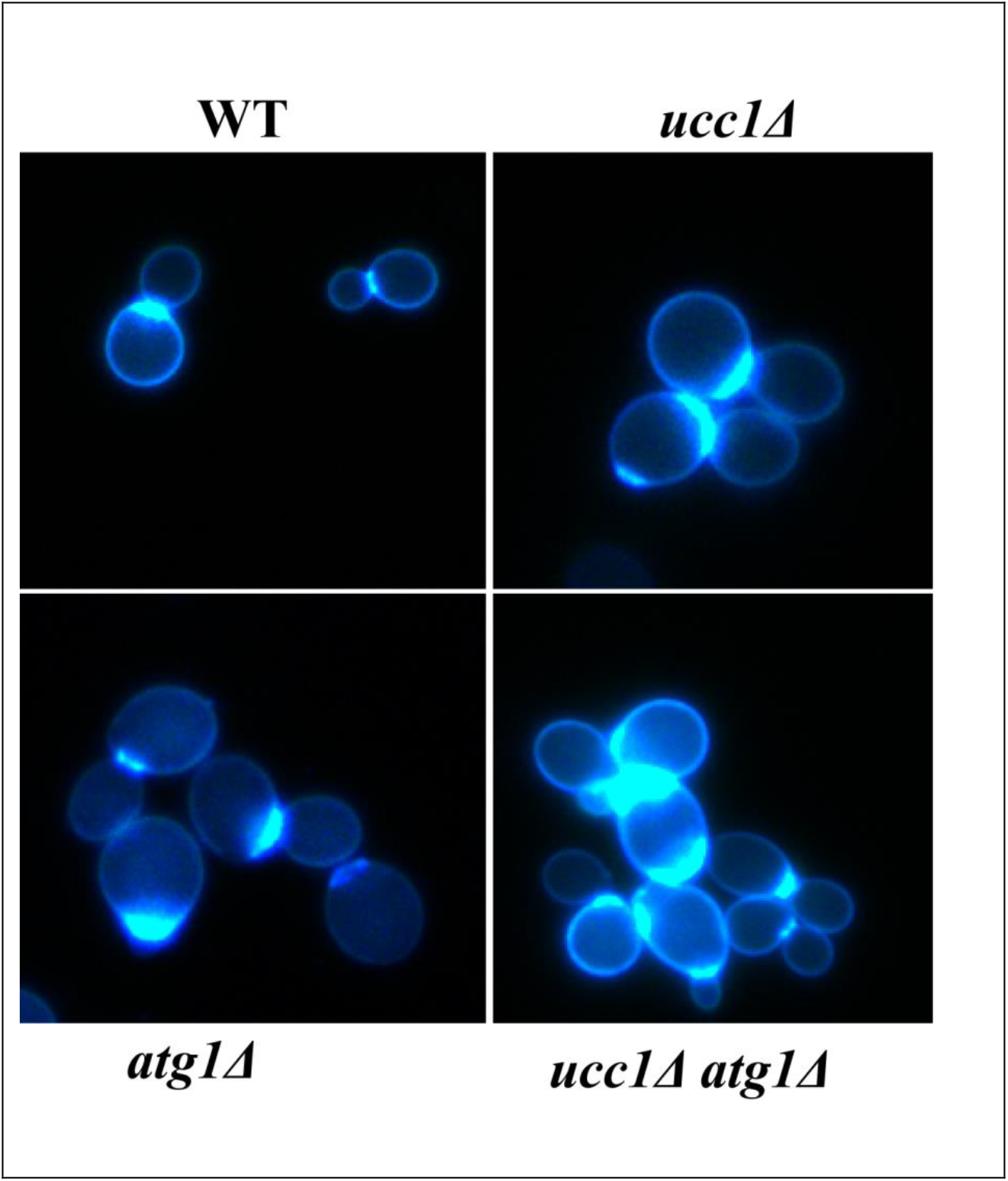
Comparative assessment of chitin distribution in WT, uc *c1Δ, atg1Δ*, and *ucc1Δatg1* Δ by Calcofluor white staining. Calcofluor white fluorescent staining was done for detection of chitin localization in cell wall. Chitin septa are visualized at the bud neck region in WT *ucc1Δ* and *atg1Δ* and *ucc1Δatg1Δ* cells. The *ucc1Δatg1Δ* cells showed the thick ring of chitin beyond neck region.

### Loss of *ATG1* and *UCC1* together leads to sensitivity to MMS, HU, and H_2_O_2_

Hydroxyurea is a non-alkylating agent and inhibitor of ribonucleotide reductase and inhibits the process of DNA replication. It also act as cytotoxic molecule and cause chromosomal damage and increased oxidative stress (Singh and Xu 2016). Hydrogen peroxide is a promoter of reactive oxygen species (ROS), oxygen radicals that are known to affect the metabolic processes and inflict damage to organelles, nucleic acids and proteins (Collinson and Dawes 1992). We studied the cellular growth response of WT, *atg1Δ, ucc1Δ* and *atg1Δucc1Δ* in presence of 0.035% MMS, 200 mM hydroxyurea, and 4mM H_2_O_2_ by spot analysis. The *atg1Δucc1Δ* cells showed the sensitivity to the stress agents (**Fig. 3a)** in comparison to WT, *atg1Δ*, and *ucc1Δ*. The morphological assessment through phase contrast microscopy of MMS and HU treated cells showed enlarged *atg1Δucc1Δ cells* when compared with WT and single mutants (**Fig. 3b)** whereas H_2_O_2_ treated *atg1Δucc1Δ* showed small size (**Fig. 2b**) when compared with WT and single mutants.

**Figure 3:**
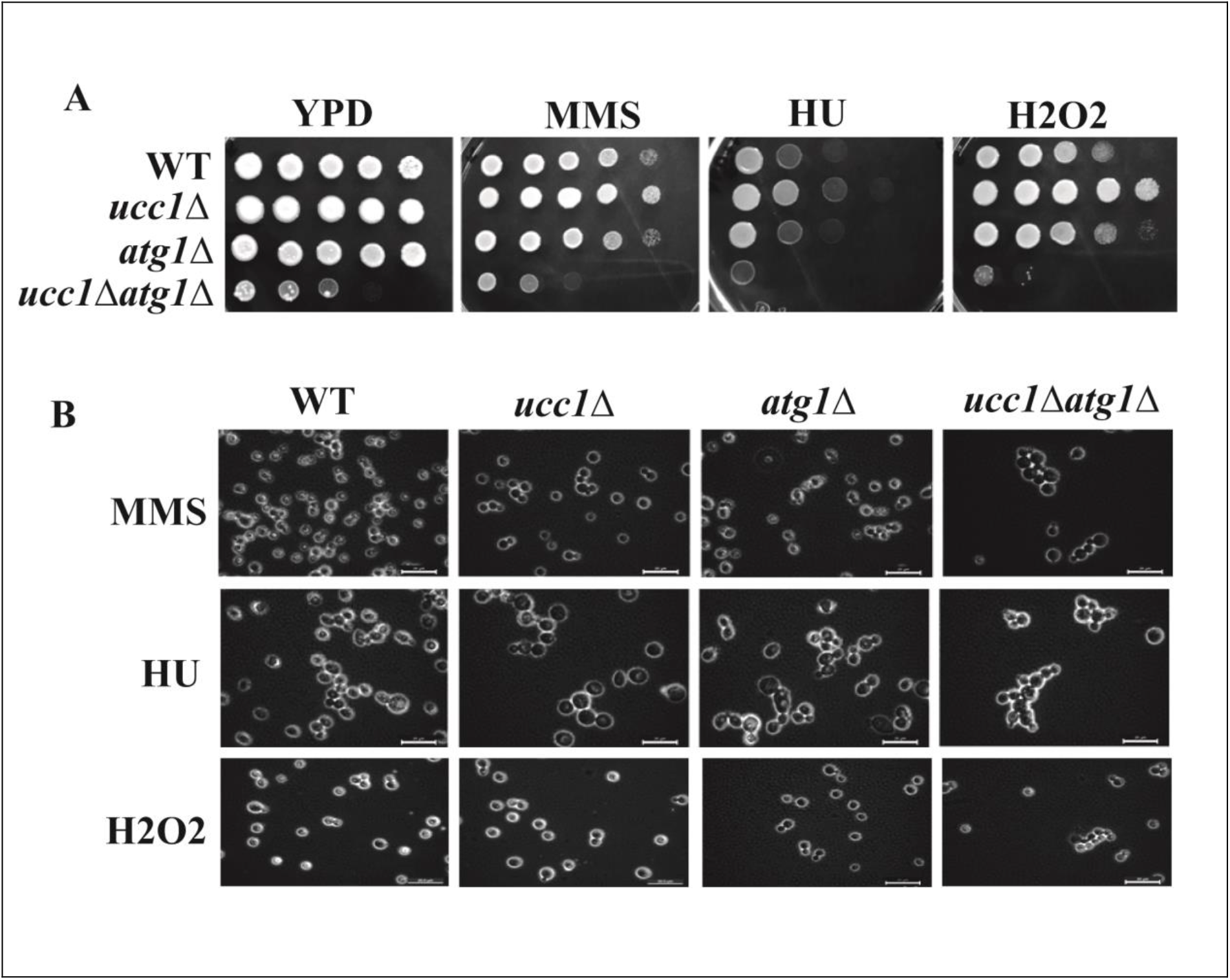
Comparative assessment of cellular growth response of *WT ucc1Δ, atg1Δ, and ucc1Δatg1Δ* cells to genotoxic and oxidative stress agents. **A.** Log phase grown cells of each strain serially diluted and spotted on YPD+ agar plate and plates containing 0.035% methyl methane sulfonate,200mM hydroxyurea, and 4mM Hydrogen Peroxide. The *ucc1*Δ*atg1*Δ showed growth sensitivity to MMS, HU and hydrogen peroxide. **B.** Comparative morphological characteristics of WT, *ucc1Δ, atg1Δ, and ucc1Δatg1Δ* cells treated with MMS, HU and H_2_O_2_ by phase contrast imaging. Cells treated with stress agents, MMS and HU indicated the increase in cell size in comparison to untreated cells.

### Absence of *ATG1* and *UCC1* together leads to altered nuclear morphology

The synthetic growth defect observed with the *atg1Δucc1Δ* cells prompted us to investigate the nuclear status of WT, *atg1Δ, and ucc1Δ* and *atg1Δucc1Δ* cells by DAPI staining. The DAPI stained *atg1Δucc1Δ* cells showed altered nuclear morphology when compared with the WT, *atg1Δ, and ucc1Δ (***Figure 4**).

**Figure 4:**
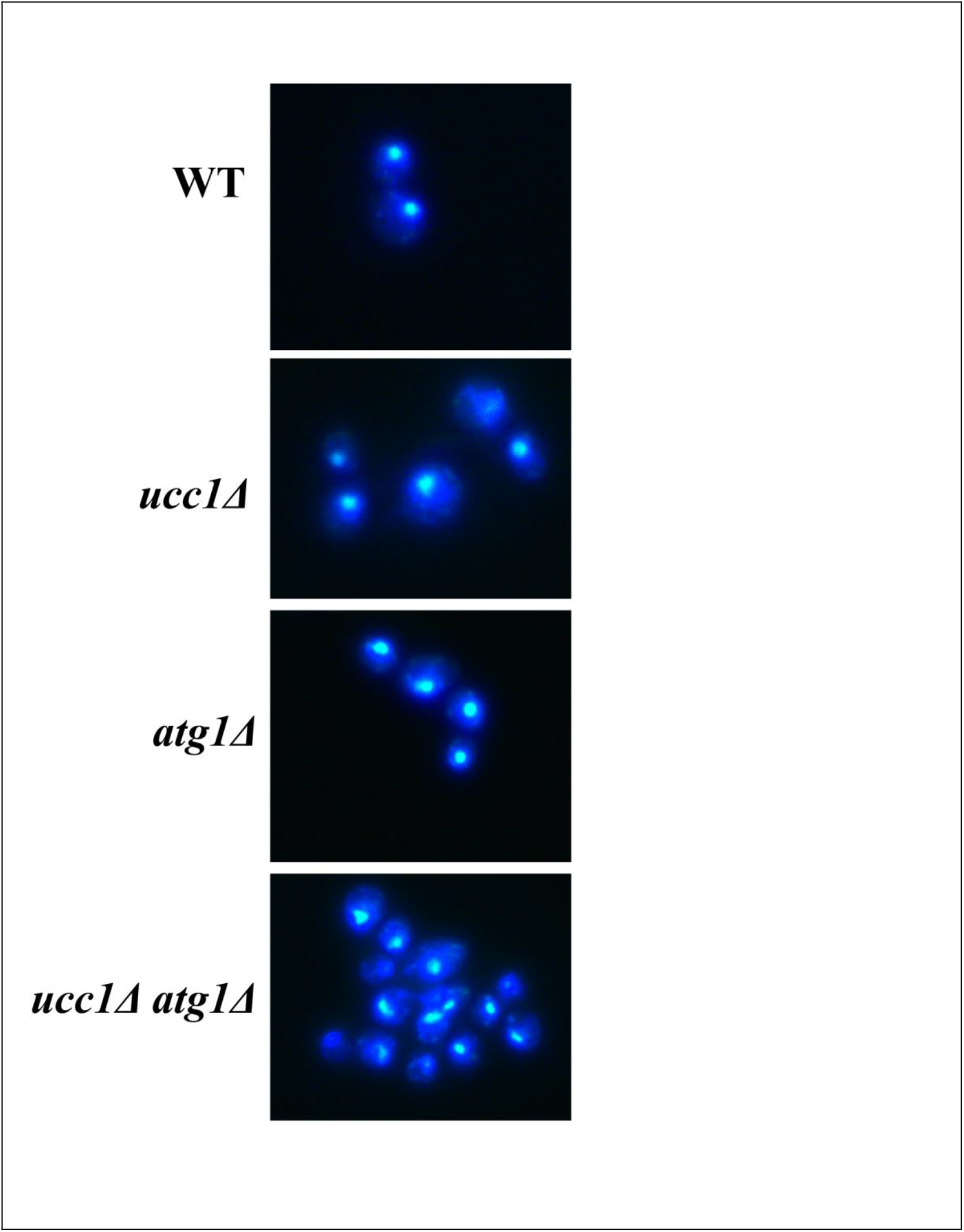
Comparative assessment of nuclear status in WT, *ucc1Δ, atg1Δ* and *ucc1Δatg1Δ* using DAPI (4′, 6-diamidino-2-phenylindole) staining. The DAPI staining revealed the altered nuclear morphology in *ucc1Δatg1Δ cells in comparison to* WT, *ucc1Δ, atg1Δ* cells.

## Discussion

Here in this study, we investigated the genetic interaction between *ATG1* and *UCC1*, representing *autophagy* and ubiquitin proteasomal system that function in the maintenance of homeostasis and cell survival. We evaluated the functional relationship by genetic interaction between the two pathways and impact on the cellular growth fitness. The *ATG1* gene product Atg1 ensures the survival of the cell under stress conditions through the autophagy whereas *UCC1* stabilizes the cellular environment by targeted proteasomal degradation of misfolded proteins (GonzÁlez-Ramos *et al.* 2013). The Ucc1 is also involved in the regulation of glyoxylate cycle by ubiquitination mediated degradation of Cit2.The loss of *ATG1* and *UCC1* together results in the synthetic growth defects. In one of our other parallel study also (Shoket et. al. 2020), we observed the synthetic growth defects phenotype with the loss of *ATG1* and F-box motif encoding gene *YDR131C with* flocculation phenotype. These studies indicated the cross talk between both the pathways in the maintenance of the cellular fitness. The loss of both the genes and increased sensitivity of the cell to the genotoxic and oxidative stress condition is interesting aspect as both the pathways inactivation resulting in the synthetic growth defects is interesting for the synthetic lethality application in the higher eukaryotes which needs to be tested further. It has been established that autophagy is critical for cell survival and exposure of autophagy deficient cells to oxidative stress results in inability to recover from the ROS accumulated cell damage, therefore cells undergo cell death (Bhatia-Kissova and Camougrand 2010). The lack of both the genes (*ATG1* and *UCC1*) results in growth sensitivity in the presence of H_2_O_2_, suggesting that cell may undergo cell death which need to be further. The mechanism of synthetic growth defects in *atg1Δucc1Δ* would be explored in the future studies.

## Acknowledgment

The authors would like to thank Dr. Deepak Sharma, IMTECH, Dr. Ravi Manjithya, JNCASR, Dr. Jitendra Thakur, NIPGR, New Delhi, India for strains and plasmids. We also thank Dr. Preeti Sharma and Mr. Parvez S. Slathia at SMVDU for allowing the use of lab equipment facility.

## Compliance with Ethical Standards

Authors declares no conflict of interest.

## Ethical Approval

This article does not contain any studies with human participants performed by any of the authors.

## Author’s contributions

NKB conceived and directed the study and wrote the paper with all the authors. MP, VS, MS, HS, SP, PK conducted the experiments. All the authors analysed the data, reviewed the results, and approved the final version of manuscript.

